# Sequential acquisition of virulence and fluoroquinolone resistance has shaped the evolution of *Escherichia coli* ST131

**DOI:** 10.1101/039123

**Authors:** Nouri L. Ben Zakour, Areej S. Alsheikh-Hussain, Melinda M. Ashcroft, Nguyen Thi Khanh Nhu, Leah W. Roberts, Mitchell Stanton-Cook, Mark A. Schembri, Scott A. Beatson

## Abstract

*Escherichia coli* ST131 is the most frequently isolated fluoroquinolone resistant (FQR) *E. coli* clone worldwide and a major cause of urinary tract and bloodstream infections. Although originally identified through its association with the CTX-M-15 extended-spectrum β-lactamase resistance gene, global genomic epidemiology studies have failed to resolve the geographical and temporal origin of the ST131 ancestor. Here, we developed a framework for the reanalysis of publicly available genomes from different sources and used this dataset to reconstruct the evolutionary steps that led to the emergence of FQR ST131. Using Bayesian estimation, we show that point mutations in chromosomal genes that confer FQR coincide with the first clinical use of fluoroquinolone in 1986, and illustrate the impact of this pivotal event in the rapid population expansion of ST131 worldwide from an apparent origin in North America. Furthermore, we identify key virulence factor acquisition events that predate the development of FQR, suggesting that the gain of virulence-associated genes followed by the tandem development of antibiotic resistance primed the successful global dissemination of ST131.

## INTRODUCTION

*Escherichia coli* sequence type 131 (ST131) is a recently emerged multidrug resistant clone associated with urinary tract and bloodstream infections. *E. coli* ST131 was originally identified due to its strong association with the CTX-M-15-type extended-spectrum-beta-lactamase (ESBL) allele (1), and is now the predominant fluoroquinolone resistant (FQR) *E. coli* clone world-wide (2-4).

ST131 belongs to subgroup 1 from *E. coli* phylogroup B2, with most strains of serotype O25b:H4 (1-4). Two previous genomic studies have explored the ST131 clonal structure (2, 5) and identified a globally dominant FQR sublineage defined as clade C (2) or H30-R (5). Two additional well-supported ST131 sublineages, referred to as clades A and B, have also been described (2). Each of these clades contain a defined marker allele for the type 1 fimbriae *fimH* adhesin; *H41* (clade A), *H22* (clade B) and H30 (clade C) (6). Further analysis of clade C/H30-R ST131 identified a smaller subset of strains containing the bla_CTX-M-15_ ESBL allele referred to as clade C2 or H30-Rx (2, 5). The ST131 strain EC958 is a reference FQR clade C strain that has been well-characterised at the genomic and phenotypic level (3, 7-12).

Several early studies demonstrated variation in the complement of virulence genes in ST131, with only a few virulence factors consistently identified in all strains (1, 4, 13-15). Our comprehensive analysis of 95 *E. coli* ST131 genomes revealed that the virulence and mobile genetic element (MGE) profile was in fact consistent with the phylogenetic structure of the ST131 lineage, with clade C strains sharing a generally conserved set of genes. In contrast, the plasmid profile of ST131 is highly disparate with multiple different replicons found in closely related strains and multiple genomic contexts for the clade C2 defining bla_CTX-M-15_ ESBL gene (16, 17).

Despite its successful dissemination globally, little information is available about evolution and emergence of ST131. Both recent independent genomic studies demonstrated that ST131 emerged from a single ancestor and that most strains belong to clade C*/H*30-R (2, 5). Notably, we found that recombination accounted for the majority of variation within the ST131 lineage and recombination events were associated with the positions of MGEs (2). However, despite the number of isolates in both studies, neither resolved the geographical or temporal origin of the ST131 ancestor. In contrast, studies of other large sets of bacteria with geographical or temporal separation have determined accurate dates of divergence of important clades using statistical analyses such as the Bayesian framework implemented in BEAST (Bayesian Evolutionary Analysis by Sampling Trees) (18). For example, Glaser identified tetracycline resistance as the major driver of diversification amongst the global population of Group B *Streptococci* (19). Similarly, a large study of Methicillin resistant *Staphylococcus aureus* (MRSA) was able to date the emergence of a FQR clade to the mid-1980’s (20). These studies motivated us to combine data sets from our geographically diverse previous study (2) and from the temporally diverse Price et al study (5) to investigate the evolution of ST131 with the highest possible resolution.

## RESULTS AND DISCUSSION

### Curation of a high-quality ST131 genome sequence dataset

We first sought to obtain a high-quality set of data to carry out our analyses. 199 draft Illumina paired-end *E. coli* ST131 genomes were retrieved from public read data repositories (Dataset S1). Initial phylogenetic analyses of *de novo* and reference-guided assemblies of all 199 *E. coli* ST131 genomes indicated that several draft genomes were of low quality. Suboptimal genome data quality could interfere in subsequent phylogenetic analyses and may invalidate conclusions drawn from tree topologies. To ensure that only high quality sequences were included in our analyses, we removed 14 genome datasets that were determined to be outliers according to at least one of our assembly or mapping metrics, including number of uncalled bases, number of scaffolds and assembled genome size (Supplementary Fig. S1). This quality control filter is broadly applicable to reanalyzing public genomic data from multiple sources.

### Phylogenomic analysis of ST131

We next carried out phylogenetic reconstruction using our combined dataset of 185 Illumina paired end sequences, which represented strains from humans (n=167), animals (n=15) or other (n=3) sources isolated from USA, Canada, New Zealand, Australia, Spain, India and UK between 1967 and 2011 (Supplementary Table 1; Fig. 1). Sequence read mapping of these 185 high-quality ST131 genomes (and simulated reads from the SE15, EC958 and JJ1886 complete genomes) to the ST131 clade C reference genome *E. coli* EC958 defined 21,373 substitution Single Nucleotide Polymorphisms (SNPs) that were used to create an unrooted Maximum Likelihood phylogeny (Supplementary Fig. 2a). An independent phylogenetic tree produced with the kSNP alignment-free method was consistent with the overall topology of the ML trees (Supplementary Fig. 3). Using a Bayesian modeling algorithm, we identified 204 non-overlapping segments encompassing 1.542 Mb and containing 15,902 substitution SNPs that were introduced into the ST131 lineage by recombination (Supplementary Fig. 4 and 5; Supplementary Table 2). The length of recombinant sequence is higher than previously reported (2) as the larger dataset increases the probability that one strain will have a recombinant fragment not encountered before. However, the proportion of SNPs introduced by recombination (74.4%) is consistent with our previous study and highlights the important role recombination has played in shaping the ST131 lineage (2). Exclusion of these recombinant SNPs from phylogenetic analyses reduced the number of SNPs to 5,471 and resulted in a tree that maintained the original overall topology, albeit with substantially reduced branch lengths and some major within clade re-clustering of strains (Supplementary Fig. 2b).

**Figure 1.**
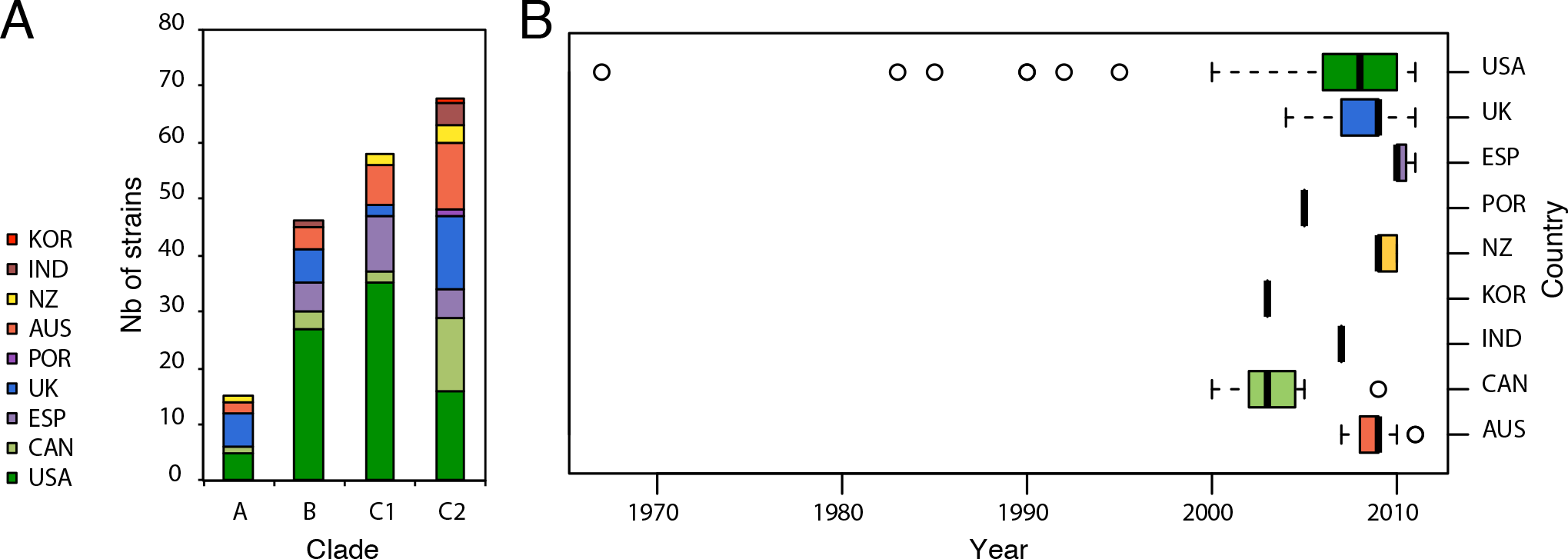
Geographical diversity of the combined dataset across clades, and time. a) Stacked histogram showing the number of strains in clades A, B, C1 and C2 according to their country of origin. The colour scheme is shown in the legend on the left along with abbreviated country names. b) Box-and-whiskers plot showing the distribution according to year of isolation for all strains based on their country of origin.

Extending our phylogenomic analyses to include isolates from two large international collections provided a far greater resolution of the evolution within the ST131 lineage (Fig. 2). The global phylogeny of *E. coli* ST131 separated the strains into three distinct lineages (clades A, B and C). Congruent with our previous work, strains in clade C were characterized by the *fimH-30* allele and the FQR conferring alleles *gyrA-1AB* and *parC-1aAB* (Fig. 2). Notable exceptions were strains JJ2643 and U004 in clade C that contain the *fimH-35* allele. This appeared to be due to a recombination event encompassing *fimH* in these strains, and highlights why we have retained a neutral nomenclature (i.e. A, B, C) for our clade classifications. Likewise, the CTX-M-15 allele is not ubiquitous in all clade C2 strains, making this a more scalable classification than the *H*30-Rx designation originally suggested by Price et al (5) (Fig. 2). In addition to harbouring CTX-M-15 genes, clade C2 strains contain more resistance genes in total compared with other ST131 clades (Fig. 3), consistent with co-localization of multiple plasmid-encoded resistance genes (Supplementary Fig. 6). Although the context of multidrug resistance cassettes can be resolved in some cases from draft genome data of ST131 isolates or transformants (16, 17), the full complexity of plasmid-mediated resistance in ST131 requires the generation of more complete genomes as per EC958 and JJ1886 (7, 21).

**Figure 2.**
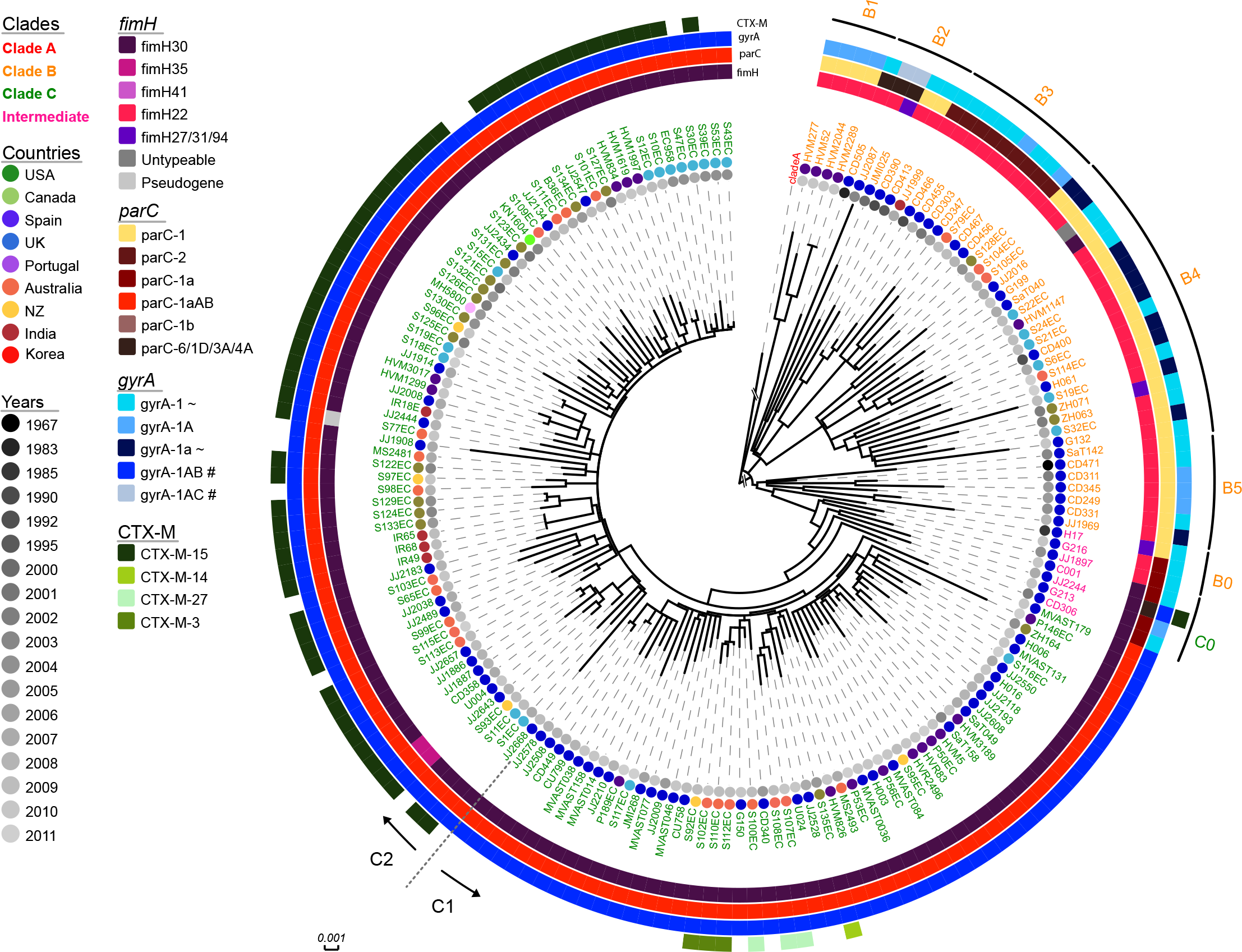
Maximum Likelihood phylogenetic tree of ST131 strains. The phylogram was built from 5,471 non-recombinant SNPs using maximum likelihood (ML). Branch support was performed by 1000 bootstrap replicates (see Supplementary Fig 2b). As shown by the scale, branch length corresponds to the number of SNP differences. Taxa labels for clades A, B and C are colored in red, orange, and green, respectively. Seven strains sharing intermediate characteristics between clades B and C are colored in pink. Of note, clade A strains were collapsed and the Clade A-specific branches shortened for display purpose. Metadata is represented as circles as follows: year of isolation in gray-gradient, and geographical region in assorted colours as depicted in the legend. Allelic profiling information is shown as coloured strips surrounding the phylogram (inner-to-outer) for *fimH, parC, gyrA,* and CTX-M. Two additional distinctions were made for some *fimH* variants: untypeable corresponds to a strain with a truncated or missing *fimH* gene; and pseudogene corresponds to a strain in which *fimH* is disrupted by an insertion sequence. Clade B0-5 and C0 sub-clades are shown as arcs in the outer-most ring with arrows and dotted lines denoting the division between sub-clade C1 and C2.

**Figure 3.**
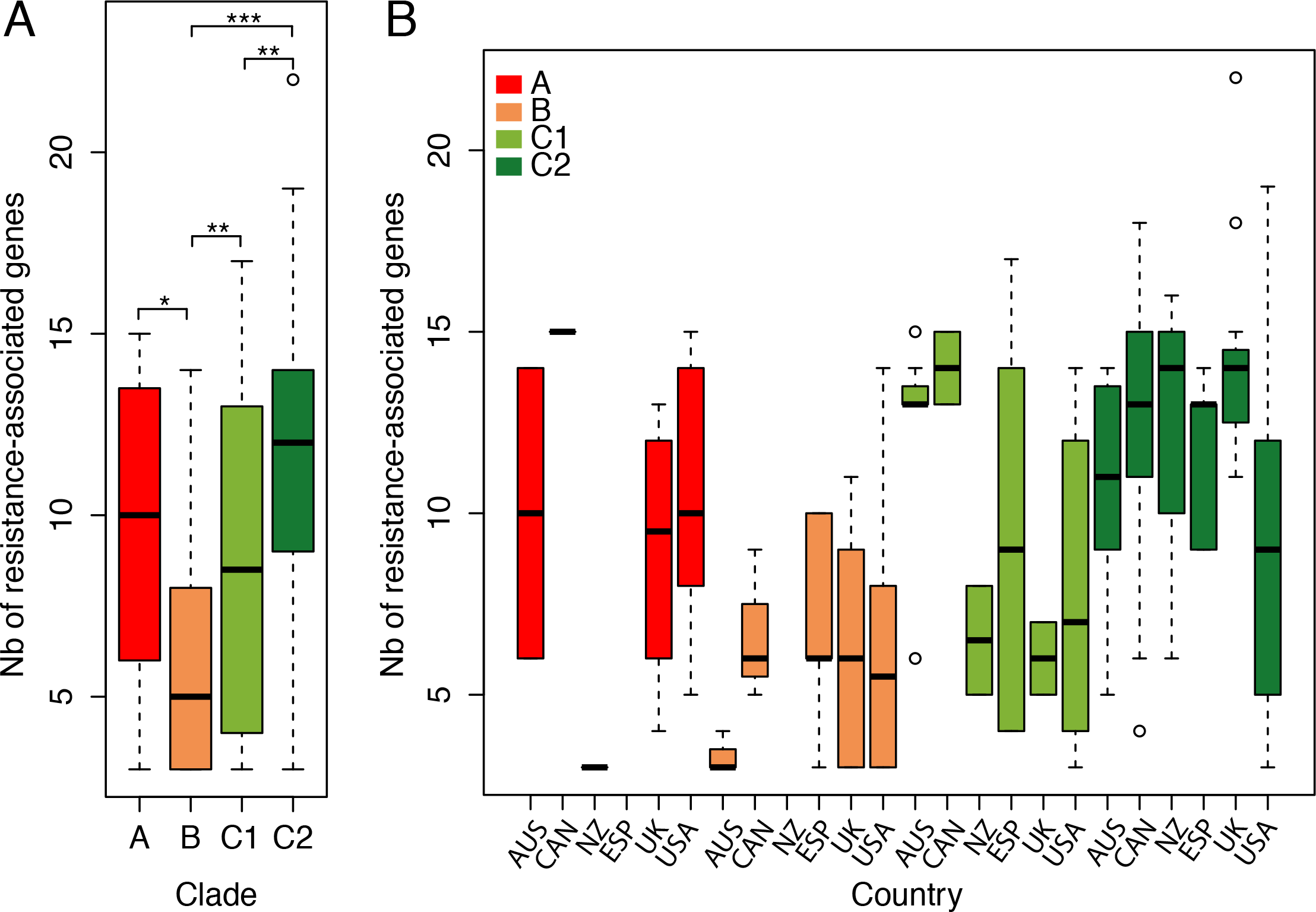
Prevalence of antibiotic resistance-associated genes by a) clade and b) clade-country. Box-and-whiskers plot showing number of resistance-associated genes per a) clade, and b) per clade and country. Colours correspond to clades, namely: clade A, red; clade B, orange; clade C1, light green; and clade C2, dark green. Screening was done using Srst2 (41) against ARGannot, with a minimum of depth of 15X read coverage. P-value are indicated as follows: ‘***’<0.001, ‘**’<0.01, ‘*’<0.05.

### Combined dataset enables greater resolution of ST131 subclades

The combined ST131 dataset enabled greater resolution of the differences between Clade B and C strains. We previously showed that Clade C strains can be further segregated into two distinct subclades, C1 and C2 (2). Our new analysis defined five discrete sub-clades in clade B (B1 to B5), each with distinct repertoires of selected marker genes *fimH, parC* and *gyrA* (Fig. 2). Notably, strains from the five clade B sublineages varied in their *parC* and *gyrA* allelic profile, with the vast majority of clade B strains containing allele combinations that are not associated with fluoroquinolone resistance. Additionally, while all strains from subclade B2 and B5 are associated with the USA, and B1 with Spain, strains within subclade B3 and B4 have a more diverse geographic origin. Each subclade showed a distinctive recombination profile (Supplementary Fig. S4 and S5) and MGE repertoire (Supplementary Fig. 7), indicative of independent evolutionary trajectories. In contrast, we found that the prevalence of virulence genes is largely conserved across all B subclades, with the absence of several UPEC-specific genes apparent in clade B3 (Supplementary Fig. 8). By comparison, our investigation of clade C MGEs and other regions of interest (as originally defined in the clade C reference strain EC958) showed a high degree of conservation across clade C, with the exception of the prophage Phi6, the capsule loci and GI*-selC* (Supplementary Fig. 7). For example, GI *selC* is only found in a geographically homogeneous cluster of clade C strains that include EC958 and excludes the reference strain JJ1886 (Supplementary Fig. S7, Fig. 2). Despite the general conservation of gene content within clade C genomes, it is apparent that genomic islands are hotspots for insertions, deletions (indels) and genome rearrangement (Supplementary Table 3). EC958 GI-*pheV* has several small indels relative to JJ1886 GI-*pheV*, and we have previously shown that the CMY-23 beta-lactamase gene that confers resistance to third generation cephalosporins has inserted within the EC958 GI-*leuX* whereas the JJ1886 GI-leuX element has a large duplication relative to EC958 (12) (Supplementary Table 3).

### Temporal analysis of ST131 identifies major divergence dates

Our initial studies of ST131 strains collected between 2001 and 2011 showed insufficient temporal depth to robustly date the emergence of clade C (2). By including 91 more strains from Price et al., (2013) (5), including 8 that predated 2000 (Fig. 1), we anticipated that we would be able to resolve this question using existing public data alone. We generated a linear regression of the genetic distance from the root-to-tip against time for the 172 ST131 isolates within clades B and C using Path-O-Gen (22). This analysis revealed a positive correlation (R^2^ = 0.3233, P < 0.0001) confirming the molecular clock-like signal (Supplementary Fig. 9). To accurately estimate the date of divergence of clade C from clade B we employed BEAST (18). BEAST analysis rejected the strict clock and favored the uncorrelated log-normal clock model in combination with a Bayesian skyline population model (Supplementary Table 4). A mutation rate of 4.39×10^−7^ SNPs per site per year (95% highest posterior density (HPD) = 3.58 − 5.23×10^−7^) was calculated, consistent with other large-scale phylogenomic analyses of the *E. coli/Shigella* lineage (6.0−10^−7^ SNPs per site per year (95% HPD = 5.2 − 6.7×10^−7^)) (23) (Supplementary Fig. S10). Based on this approach we could estimate the divergence of the last common ancestor of clade B and C strains to have occurred between 1930 and 1958 (Fig. 4A), consistent with the Path-O-Gen prediction (Supplementary Fig. S9a). We could estimate the divergence of clade C from clade B to have occurred in 1980 (95% HPD 1973-1986), which was slightly earlier than the Path-O-Gen prediction (Supplementary Fig. S9a). Importantly, we identified that further diversification of clade C2 from clade C1 dated to 1987 (95% HPD 1983-1992), subsequent to all clade C1 and C2 strains acquiring *gyrA-* 1AB and *parC-* 1aAB alleles imparting elevated FQR (Fig. 4a). Bayesian skyline plots show a relatively constant population size over several decades, followed by a short recent expansion occurring in the late 1990’s and early 2000’s and subsequent stabilization (Fig. 2c). Interestingly, this pattern is consistent with the introduction of FQ for clinical use in 1986 (24) and the subsequent stabilization may reflect the improved stewardship of FQ (or its removal from general use). A similar phenomenon was observed for FQR amongst the MRSA ST22 global phylogeny (20). Remarkably, an identical date was identified in a recent pre-print report using 81 genomes from Price et al. (5), supplemented with ∽100 newly sequenced genomes from more geographically dispersed strains (17), highlighting the value of careful analysis of existing datasets. Although acquisition of the CTX-M-15 gene within clade C2 may be a major factor in the diversification of C1 and C2, it is worth emphasizing that this alone does not explain the success of ST131 given that the population expansion identified in our study encompasses both clade C1 and clade C2 strains.

**Figure 4.**
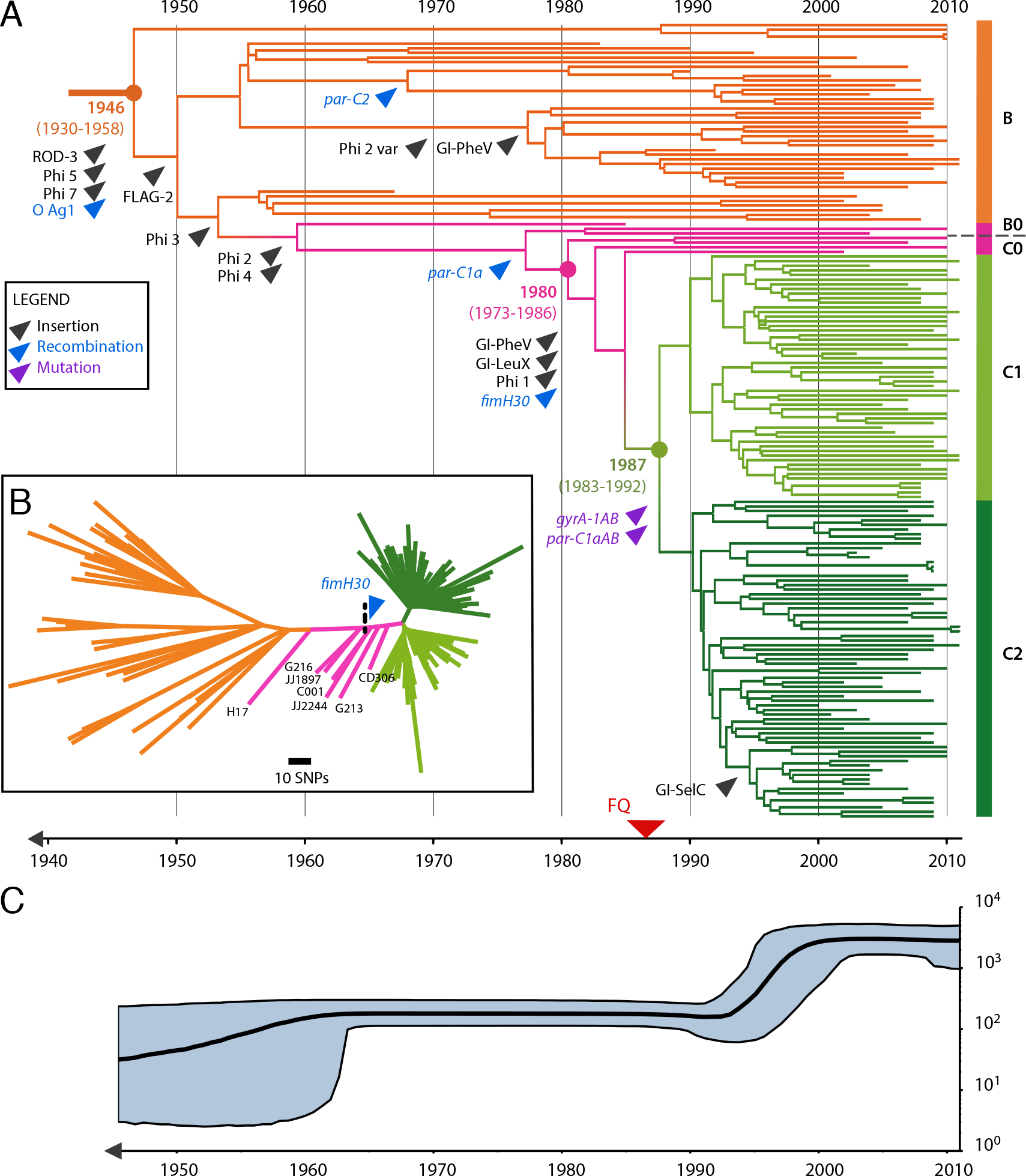
Evolutionary scenario of the emergence of ST131 clades B and C. A time-calibrated phylogeny was reconstructed using BEAST 2.0 based on 3,779 bp non-recombinant SNPs for the 172 clade B and C strains. Of all combinations tested (Table S4), the one combining the GTR substitution model, a constant relaxed clock model and the Bayesian Skyline population tree model was preferred. a) Maximum Clade Credibility tree colored according to clade origin as shown on the right with B, orange; Intermediate B(0) and C(0), in pink; C1, in light green; and C2, in dark green. X-axis indicates emergence time estimates of the corresponding strains. Major evolutionary events were also indicated by an arrow pointing at the branch onto which they are predicted to have occurred (position along the branch is arbitrary). Three categories of major events are displayed, namely: MGE and genomic island insertion events, in black; allelic change acquired through recombination events, in blue; and allelic change acquired through point mutation events, in purple. Of note, the two point mutations indicated by a purple arrow and pointing at the branch from which clade C1 and C2 originate, confer resistance to fluoroquinolone, for which the first introduction is indicated in the bottom timeline by a red arrow. b) Unrooted phylogenetic tree built on the same 3,779 bp non-recombinant set of SNPs using maximum likelihood (ML). Branch support was performed by 1000 bootstrap replicates. Intermediate strain names and predicted acquisition of *fimH30* are indicated on the tree. C) The Bayesian Skyline plot illustrates the predicted demographic changes of the ST131 clade B and C population since the mid 1940’s. The black curve indicates the effective population size (VVe) with 95% confidence intervals shown in grey.

### Phylogeography of ST131

To investigate the geographical context underlining the expansion of the multidrug-resistant ST131 O25b:H4 clone from clade B to C, we performed a discrete phylogeographical analysis as implemented in BEAST on the 172 ST131 isolates within clades B and C that included a variety of geographic sources (Fig. 1a) and dates (Fig. 1b). To reduce sampling biases due to the high number of strains isolated from USA, Canada, UK and Spain, we performed independent analyses on 10 randomly subsampled datasets containing 85 strains each (Supplementary Table 5). Under a BSSVS and symmetric diffusion models, results systematically supported the USA (74.31%, s.d. 12.1) as the most likely origin of clade B and C (Fig. 5a). The origin of clade C (C0, C1 and C2) was predicted to be associated either with USA (51.83%, s.d. 35.5) or Canada (45.59%, s.d. 36.1), overall North America (Fig. 5b). These results are consistent with the observation that the oldest reported ST131 strain was isolated in the USA in 1967 ^5^, and an independent BEAST analysis using a partly overlapping dataset (17). Although our resampling approach has minimized bias in strain origin, a dataset with greater diversity of strains from different geographical regions and pre-2000 isolation dates would be necessary to rule out a different origin (e.g. current datasets are under-represented in South America, Africa and many European and Asian countries). A greater number of strains would also help identify local outbreaks or clusters: with the exception of the GI*-selC* carrying clade C strains from the UK which cluster phylogenetically (Fig. 2), we did not observe other significant geographic clustering using this dataset alone.

**Figure 5.**
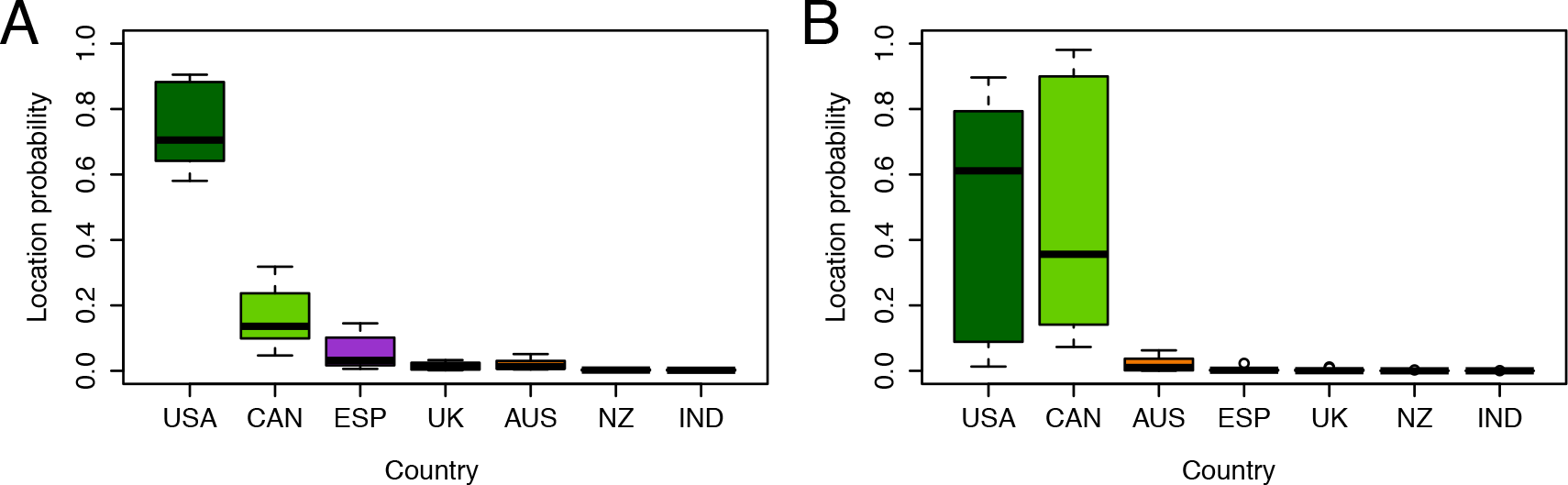
Geographic location of the Most Recent Common Ancestor (MRCA) of ST131 major clades. Individual probabilities were predicted from 10 independent BEAST analyses of randomly subsampled data to limit bias related to over-representation of some locations. Mean probability of the geographic location of the MRCA for a) clades B and C, and b) clade C, are shown as a box-and-whiskers plot colored according to country using the scheme as described in Figure 1. Countries are labeled on the x-axis by abbreviation.

### Intermediate strains reveal key MGE acquisitions that define clade C

Overall, excluding intermediate B0 and C0 strains, clade C differs from clade B by only 42 substitution SNPs (Supplementary Table 6). This list included the majority of the 70 clade C defining SNPs reported in our earlier study (2), but was not identical due to the greater number of recombinant regions identified and removed in the present study. Intriguingly, we found that seven strains originating from the USA could be classified as “intermediate” on the basis of their SNP pattern (Supplementary Table 6). These strains showed progressive acquisition of clade C-defining point mutations, with the three isolates closer to clade B (classified B0) and four isolates closer to clade C (classified C0) illustrating the precise evolutionary events leading to the emergence of clade C (Fig. 2b, Supplementary Table 5). Closer examination of the recombination analysis identified intermediate patterns of recombination, primarily clustered around known MGEs, indicative of step-wise evolution among these intermediate strains (Supplementary Fig. S4 and S5). Most notably, we could trace the acquisition of Gl*-pheV* and Gl*-leuX* genomic islands to the most recent common ancestor of the C0 strains (C001, JJ2244, G213 and CD306), several years before the acquisition of the FQR mutations that define clade C (Fig. 4). The *pheV* genomic island acquired by clade C ST131 strains is known to carry the autotransporter genes *agn43* and *sat,* the ferric aerobactin biosynthesis gene cluster *(iucA-D)* and its cognate ferrisiderophore receptor gene *iutA* (3, 7). The clade-defining *fimH30* allele was acquired by recombination (25), possibly in conjunction with the acquisition of the nearby *Gl-leuX* island; the same time-point is also predicted for the acquisition of the lS*Ec55* insertion sequence within the *fimB* recombinase gene that we have previously shown to affect the expression of type 1 fimbriae (3, 7). Thus, close scrutiny of these “intermediate” genomes enabled us to trace the acquisition of virulence-associated genes in ST131, which appears to have primed this clone for success prior to the acquisition of FQR mutations in the late 1980’s. Further molecular analysis is required to determine the contribution of these elements to ST131 colonization/fitness in the gastrointestinal and urinary tracts. Notably, the role of virulence in the success of this clone may have been underappreciated in a recent report as these particular strains were distributed throughout clade B and clade C despite their inconsistent *parC, gyrA* and *fimH* alleles (17).

## CONCLUSIONS

Overall, our work highlights how careful reanalysis of publicly available genomic datasets from heterogeneous sources can greatly improve the resolution of evolutionary history. Here, we have characterised the evolution of ST131 with unprecedented detail, from the acquisition of prophages and the O25b antigen region circa 1946, to the acquisition of GI-*pheV*, GI-*leuX*, *fimH30* and lS*Ec55* around 1980, several years before the acquisition of mutations in *gyrA* and *parC* that led to FQR and the acquisition of the clade C2 defining CTX-M-15 ESBL gene. Whereas the development of FQR was accompanied by a large surge in the ST131 population globally, we propose that the acquisition of virulence factors by ST131 was a necessary precursor to this success. These events describe the ‘perfect storm’ for the evolution of a multidrug resistant pathogen; the acquisition of virulence-associated genes followed by the development of antibiotic resistance.

## MATERIALS AND METHODS

### Genome data

Two *E. coli* ST131 strain datasets from previously published work were used in this study, under the designation Dataset_1 and Dataset_2 (2, 5). Strain names, sources and available strain metadata are summarized in Supplementary Table 1. Dataset_1 comprised Illumina 101-basepair (bp) paired-end genome sequence data of 95 ST131 strains isolated from 2000 to 2011, mostly in Europe and Oceania (2) (study accessions ERP001354 and ERP004358). Dataset_2 comprised Illumina 101-bp (76 samples), 76-bp (19 samples) and 50-bp (10 samples) paired-end whole genome sequence data of 105 ST131 strains isolated from 1967 to 2011, mostly in North America (5) (study accession SRP027327). Additionally, reference strains of 11 published complete genomes were also used, namely *E. coli* ST131 strains SE15, JJ1886 and EC958, plus non-ST131 B2 phylogenetic group *E. coli* strains CFT073, UTI89, E2348, ED1a, 536, S88, APEC-01 and non-ST131 D phylogenetic group *E. coli* strain UMN026 (Supplementary Table 7). *E. coli* strain NA114 was excluded from the analysis due to poor assembly quality (2, 7, 26). To integrate reference genome data into our phylogenomic analyses, error-free 101-bp paired-end Illumina reads were simulated to 60 times coverage with an insert size of 340 bp +/− 40 bp as previously described (2).

### Quality control, de novo genome assembly and variant detection

Quality Control (QC) was performed for all raw read datasets. Briefly, raw reads were analyzed using PRINSEQ v0.20.3 (27) and trimmed with a mean base-pair quality score Q ≥ 20 and a read length of ≥ 70% of the original read length. Additionally, it was necessary to correct 35 sets of raw read data from Dataset_2 that had heterogeneous Illumina encoding and/or erroneous paired-end length encoding (Supplementary Table 8). Quality control and assembly metrics for Dataset_1 have been previously reported in Petty et al. (2). Lastly, contaminant searches were performed for each sample using Kraken on a subset of 100,000 randomly chosen reads (28).

Quality filtered Illumina paired-end reads were assembled *de novo* using Velvet v1.2.07 (29) with a &-mer range of 45-85 for 101-bp reads, 29-61 for 76-bp reads, and 29-47 for 50-bp reads. An optimal *k*-mer value for each assembly was selected on the basis of best assembly metrics including N50 (50% of bases are incorporated in contigs of this length or above), number and size of contigs, number and continuity of uncalled bases and peak coverage. Contigs ≥ 200 bp at an optimal *k*-mer were then ordered against *E. coli* EC958 (7) using Mauve v2.3.1 (30). All QC and assembly statistics are summarized in Supplementary Table 8.

Quality filtered Illumina reads for Dataset_1 and Dataset_2, as well as error-free simulated reads of complete genomes, were mapped on the reference strain EC958 using SHRIMP v2.0 (31). Nesoni v0.108 (32) was used to call and annotate substitution-only SNPs, with a consensus cutoff and majority cutoff of 0.90 and 0.70, respectively. SNPs were also determined in parallel using the reference-free *k*-mer based approach developed in kSNP v2.0 (33). Default parameters as well as a *k*-mer value of 19 selected as the optimal value predicted by the kSNP associated Kchooser script were applied.

### Exclusion of suboptimal genome datasets

We devised a statistical approach that excluded outliers based on several non-Gaussian metrics that could be determined from mapped and assembled Illumina genome data (summarized in Supplementary Table 8). Specifically we examined three mapping metrics: (i) sequence coverage, (ii) number of unmapped bases (in the mapping reference EC958 genome) and (iii) number of uncalled bases due to low coverage or mixed-base calls); and two assembly metrics: (iv) the number of scaffolds >= 200bp and (v) estimated genome size. Sub-optimal genomes were discriminated quantitatively on the basis of metrics (iii), (iv) and (v), and a total of 14 outliers were identified (based on upper and lower cut-offs at the Quartile 3 + 1.5 Interquartile range and the Quartile 1 - 1.5 Interquartile range cut-offs, respectively). Metric (i) did not identify any outliers with low sequence coverage and outliers with high sequence coverage were not omitted, whereas metric (ii) did not discriminate any outliers. This additional QC process resulted in the exclusion of genome data from the following strains CD301, CD436, JJ1901, JJ1996, JJ2007, JJ2041, JJ2050, JJ2243, JJ2441, JJ2555, MH17102, QU300, QUC02 and ZH193. R scripts used are available on github at https://github.com/BeatsonLab-MicrobialGenomics/ST131-200. A final dataset of 188 ST131 genomes (including the complete genomes of EC958, JJ1886 and SE15) were chosen for further study after excluding the 14 genomes with suboptimal data quality.

### Recombination detection

To avoid distortion of the phylogenetic signal caused by SNPs acquired through recombination, we used the Bayesian clustering algorithm BRATNextGen (34) to detect recombinant regions among the combined dataset. Similar to our previous work (2), we used as an input a SNP-based multiple genome alignment composed of each strain-specific pseudo-genome built by integrating the SNPs predicted for each strain to the reference genome of EC958. To help identification of underlying clusters of strains, BRATNextGen initially computes a hierarchical clustering tree relative to the proportion of ancestral sequences shared between all strains. A segregation cut-off of 0.12 was then specified to separate each previously identified ST131 clade (clades A, B and C) and non-ST131 strains into distinct clusters. Recombination was then evaluated within and between each cluster with the convergence approximated using 20 iterations of the learning algorithm. Significance was estimated using 100 permutations with a statistical significance threshold of 0.05.

### Phylogenetic analysis

SNPs identified through reference-based mapping for the 188 ST131 strains were used to build phylogenies using Maximum Likelihood (ML), prior to and after removal of SNPs associated with recombinant regions. Phylogenetic trees were generated with RAxML v7.2.8 (35) using the general time reversible (GTR) GAMMA model of among site rate variation (ASRV), and validated using 1000 bootstrap repetitions to assess nodal support. Additionally, reference-free *k*-mer based phylogenetic trees were constructed using kSNP v2.0 with default parameters (33) and genome assemblies as an input. A *k*-mer value of 19 was selected as the optimal value predicted by the kSNP associated Kchooser script. All trees were then viewed using Figtree v1.4.0 (36) or Evolview (37), and further compared using the Tanglegram algorithm of Dendroscope v3.2.10 (38), which generates two rectangular phylograms to allow comparison of bifurcating trees.

### Bayesian temporal and geographical analysis

Preliminary estimation of the underlying temporal signal of our data was obtained by performing a regression analysis between the root-to-tip genetic distance extracted from the recombination-free maximum-likelihood tree, the isolation year and lineage information for each sequence, as implemented in Path-O-Gen v1.4 (22). To further investigate the divergence of clade C from clade B, we performed a temporal analysis on the 3,779 bp non-recombinant SNPs of the 172 clade B and C strains using BEAST 2.0 (18), a Bayesian phylogenetic inference software, which can estimate the dating of emergence of distinct lineages. We compared multiple combinations of molecular clock model (strict, constant relaxed lognormal, and exponential relaxed lognormal), substitution model (HKY, GTR) and population size change model (coalescent constant, exponential growth, Bayesian skyline, extended Bayesian skyline). Markov Chain Monte Carlo generations for each analysis were conducted in triplicate for 100 million steps, sampling every 1,000 steps, to ensure convergence. Replicate analyses were then combined with LogCombiner, with a 10% burn-in. The GTR nucleotide substitution model was preferred over the HKY model, and was used with four discrete gamma-distributed rate categories and a default gamma prior distribution of 1. The uncorrelated lognormal clock model consistently gave better support based on the Bayes Factor and AICM analyses, compared to a strict clock model. The Bayesian skyline population tree model was chosen as the best fitting tree model. Maximum clade credibility (MCC) trees reporting mean values with a posterior probability limit set at 0.5 were then created using TreeAnnotator.

In order to adequately investigate the biogeographical history of our ST131 collection, we evaluated potential bias in the geographical origin of strains, which could negatively impact our predictions. Statistical significance of the geographical origin distribution in clade B, C1 and C2 was assessed by Chi-Square test with Bonferroni correction for multiple comparisons. Over-represented countries were randomly subsampled down to 15 representative sequences, while countries with fewer than 5 representatives had to be excluded from the analysis (Korea and Portugal). Overall, we constructed 10 independent randomly subsampled datasets with 85 isolates representing 7 countries, each with 5 to 15 representative sequences. Reconstruction of possible ancestral geographical states was then performed using BEAST 1.8.2 on each individual subsampled dataset. In addition to the previous parameters selected for the temporal analysis, a symmetric substitution model, a Bayesian stochastic search variable selection (BSSVS) model and a strict clock for discrete locations were chosen for the phylogeographical analysis. MCMC generations were conducted for 100,000,000 steps, sampling every 10,000 steps. MCC trees were then generated using TreeAnnotator for each run with a posterior probability limit set at 0.5. Location posterior probabilities of MRCA were then collated for clade B and C, and clade C only.

### Genomic comparisons and in silico genotyping

Comparative genomic analyses were conducted using a combination of tools, namely Artemis, Artemis Comparison Tool (39) and Mauve (30). Graphical representations showing the presence, absence or variation of mobile genetic elements (MGE) or other regions of interest, virulence factor genes, and antibiotic resistance genes were carried out using BLASTn and read-mapping information as implemented in the Seqfindr visualization tool (40). Regions of interest previously described in the genome of ST131 reference strain EC958 (2, 7) and virulence factors including autotransporters, fimbriae, iron uptake, toxins, UPEC-specific genes and other virulence genes were screened in all ST131 strains with SeqFindr using a cut-off >= 95% nucleotide identity over the whole length when compared to the assembly or the consensus generated from mapping. Additionally, prevalence of antibiotic resistance-associated genes was also investigated using Srst2 (41) against ARGannot database, with a minimum of depth of 15X read coverage.

